# cgCorrect: A method to correct for confounding cell-cell variation due to cell growth in single-cell transcriptomics

**DOI:** 10.1101/057463

**Authors:** Thomas Blasi, Florian Buettner, Michael K. Strasser, Carsten Marr, Fabian J. Theis

## Abstract

**Motivation:** Accessing gene expression at the single cell level has unraveled often large heterogeneity among seemingly homogeneous cells, which remained obscured in traditional population based approaches. The computational analysis of single-cell transcriptomics data, however, still imposes unresolved challenges with respect to normalization, visualization and modeling the data. One such issue are differences in cell size, which introduce additional variability into the data, for which appropriate normalization techniques are needed. Otherwise, these differences in cell size may obscure genuine heterogeneities among cell populations and lead to overdispersed steady-state distributions of mRNA transcript numbers.

**Results:** We present cgCorrect, a statistical framework to correct for differences in cell size that are due to cell growth in single-cell transcriptomics data. We derive the probability for the cell growth corrected mRNA transcript number given the measured, cell size dependent mRNA transcript number, based on the assumption that the average number of transcripts in a cell increases proportional to the cell’s volume during cell cycle. cgCorrect can be used for both data normalization, and to analyze steady-state distributions used to infer the gene expression mechanism. We demonstrate its applicability on both simulated data and single-cell quantitative real-time PCR data from mouse blood stem and progenitor cells. We show that correcting for differences in cell size affects the interpretation of the data obtained by typically performed computational analysis.

**Availability:** A Matlab implementation of cgCorrect is available at http://icb.helmholtz-muenchen.de/cgCorrect

**Supplementary information:** Supplementary information are available online. The simulated data set is available at http://icb.helmholtz-muenchen.de/cgCorrect

## 1 Introduction

Recent technical advances allow for the analysis of single cells with high throughput omics technologies [Wang *et al*., 2010]. Especially single-cell transcriptome analysis [Tang *et al*., 2011, Wu *et al*., 2014] has made dramatic advances. Investigating transcripts of single cells with both quantitative real-time PCR (qPCR) [Stahlberg *et al*., 2010, Citri *et al*., 2012], and single-cell RNA sequencing (RNA-seq) [Tang *et al*., 2009, Islam *et al*., 2011, Yan *et al*., 2013, Islam *et al*., 2014] has become possible. But new experimental methods bring new challenges with them: Biological variability among single cells, which remained hidden in population based approaches now becomes evident. One major challenge of computational biology is the development of new and the adaptation of existing methods for single-cell gene expression data [Kim *et al*., 2013, Buettner *et al*., 2012].

Gene expression is a stochastic process [Elowitz *et al*., 2002] and the abundance of mRNA transcripts (of an individual gene) among many single cells (of the same cell type) can be formulated in terms of steady-state probability distributions [Thattai *et al*., 2001, Raj *et al*., 2006]. Analyzing these steady-state probability distribution can yield new insights into the underlying gene expression mechanism [Shahrezaei *et al*., 2008, Larson, 2011, Kim *et al*., 2013].

There are two well-studied mechanisms of gene expression that have been serving as a paradigm [Raj *et al*., 2008]: Simple, constitutive gene expression (also known as birth-death process), where DNA is continuously transcribed to mRNA (see Figure 1A) and bursty gene expression, where the promoter of the DNA successively switches between an active and inactive state and transcripts are produced in episodical bursts (see Figure 1B). The steady-state distributions of simple gene expression follows the Poisson distribution [Peccoud *et al*., 1995, Thattai *et al*., 2001] whereas the steady-state distribution of bursty gene expression follows the negative binomial distribution [Raj *et al*., 2006], which allows for more variability among the transcript numbers.

Besides the stochastic nature of gene expression that gives rise to this insightful biological variability, there are also other, confounding sources of variability, such as technical noise [Ramsköld *et al*., 2012, Brennecke *et al*., 2013, Buettner *et al*., 2014, Vallejos *et al*., 2015] and cell cycle effects. Especially the influence of the latter on the interpretation of gene expression data based on steady-state probability distributions has not been investigated so far, even though confounding cell cycle effects appear in all proliferating cells (such as stem and progenitor cells). During cell cycle, the cell grows and the number of transcripts within a cell doubles on average [Mitichison, 2003]. Recently [Padovan-Merhar *et al*., 2015] found experimental evidence for the compensation of differences in cell size and suggest that the concentration of transcripts within a cell is maintained constant. This means that measuring the abundance of a particular transcript in two identical cells with different cell sizes will yield different results. The differences in cell size cause a broadened, overdispersed steady-state distribution of transcript numbers, which may be mistakenly interpreted in an upstream analysis.

To illustrate this issue we consider the following scenario (illustrated in Figure 1C): Assume we measure the mRNA transcripts of a particular gene from several single cells, which have the same volume. The gene of interest is subjected to simple, constitutive gene expression and follows the Poisson distribution. In a typical experiment, however, cells are not synchronized and single cells with different sizes are pooled together (see Figure 1C) leading to an overdispersed steady-state distribution. Performing model selection (see Methods) on the steady-state distribution of transcript numbers obtained by this type of experiment incorrectly favors the negative binomial over the Poisson distribution and therefore the gene expression mechanism would be interpreted to be bursty.

Here, we introduce cgCorrect (cell growth correction), a statistical method to correct single-cell transcriptomics data for latent differences in cell size. cgCorrect can be used for both normalizing single-cell gene expression data sets, and for parameter estimation and model selection on steady-state distributions of gene expression. Our approach is based on the assumption that the average number of mRNA transcripts within the cell increases proportionally to the volume as the cell grows during cell cycle, leaving the concentration of transcripts constant [Padovan-Merhar *et al*., 2015].

We calculate the cell growth correction probability, which corrects for differences in transcript numbers that are due to differences in cell size. This is the conditional probability, for finding the corrected, cell growth independent number of mRNA transcripts of a particular gene, given the measured, cell growth dependent number of mRNA transcripts of this gene. cgCorrect can include information on the cells’ volume, but, more strikingly, it can also be applied if there is a total lack of additional information on the cell’s volume. Since the cell volume is typically not observed, we marginalize this latent variable out, which corresponds to a blind deconvolution problem.

cgCorrect is based on discrete molecule numbers of individual mRNA transcripts in single cells. Discrete molecule numbers are essential for the interpretation of the underlying mechanism of gene expression [Raj *et al*., 2009]. There are two high throughput transcriptomics techniques, qPCR and RNA-seq, which both hold the ability to measure discrete molecule numbers in single cells (e.g. via digital PCR [Vogelstein *et al*., 1999], droplet digital PCR [Hindson *et al*., 2011], direct RNA sequencing [Ozsolak *et al*., 2009] or strand-specific single-cell sequencing [Islam *et al*., 2011]). If the experiment does not provide discrete molecule numbers, the data can be converted to such by matching the measured value (e.g. cycle time (ct) values in qPCR experiments) to known absolute molecule numbers of a particular gene in the same cell type.

**Fig. 1.**
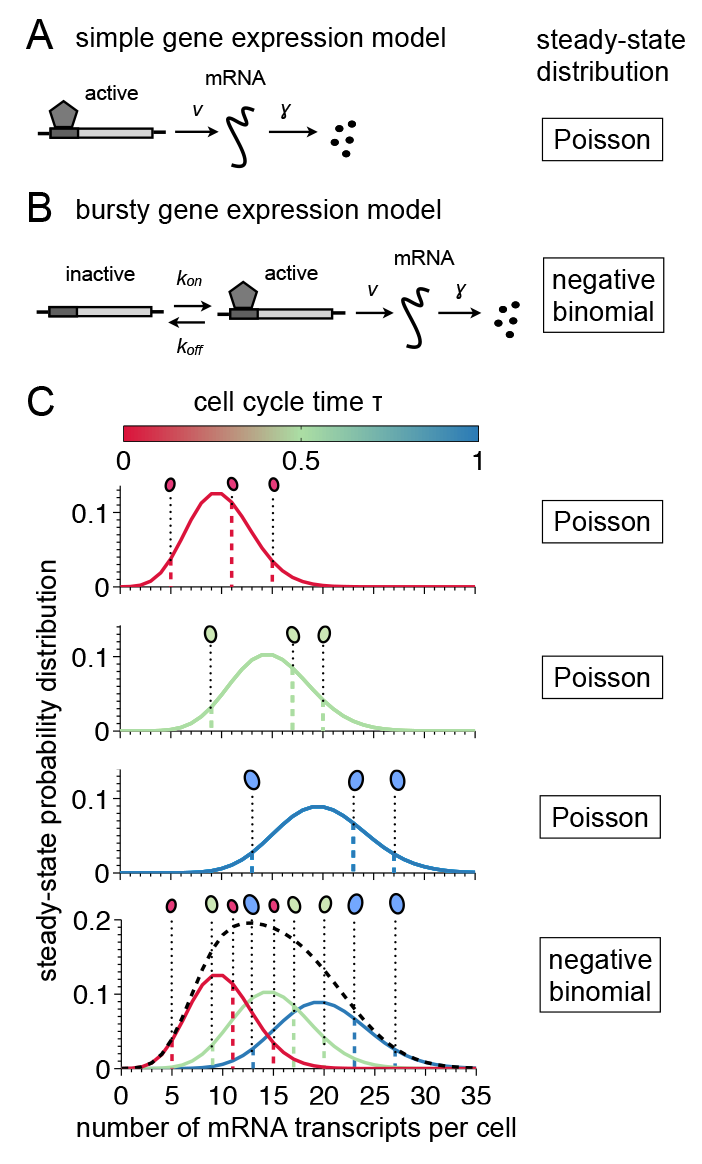
Differences in cell size lead to an overdispersed mRNA distribution. (A) Simple gene expression mechanism. The promoter of a particular gene is always active transcribing its associated DNA to mRNA with rate *ν*. The degradation rate of the mRNA is given by *γ*. (B) Bursty gene expression mechanism. Additionally to the simple gene expression mechanism, the promoter can perform transitions between the active and inactive state with rates *k*_on_ and *k*_off_, respectively. (C) A measurement of mRNA transcripts from single cells with different cell sizes that are pooled together leads to an overdispersed steady-state distribution. We display the Poisson distribution of steady-state transcript numbers for three generic cells with different cell sizes (increasing volume from top to bottom) that are all subjected to constitutive, simple gene expression. By pooling these 9 cells together and ignoring their different volumes we obtain an overdispersed steady-state distribution of transcript numbers (bottom panel, dashed black line) that does not follow a Poisson distribution anymore. The overdispersed distribution may be mistakenly interpreted to be due to the bursty gene expression mechanism.

Current state-of-the-art normalization techniques to account for confounding variability are based on scaling the measured number of mRNA transcripts with reporters that should correlate with the confounding variability. In qPCR where the mRNA transcripts of only a few genes are observed the measured number of transcripts is scaled with the abundance of house-keeping gene transcripts from the same single cell [Moignard *et al*., 2013, Guo *et al*., 2010, Liviak *et al*., 2013]. In RNA-seq experiments where the whole transcriptome is measured the sum of all mRNA transcripts or rank statistics thereof can be used as an estimator for the cell size of each single cell [Glusman, 2013 *et al*., Brennecke *et al*., 2013, Sasagawa *et al*., 2013, Vallejos *et al*., 2015]. However, scaling does not account for the discreteness of mRNA numbers.

Scaling normalization strategies can also be performed based on genes selected from the data as has been pointed out for bulk measurements [Glusman, 2013 *et al*.]. Whereas this approach is infeasible for single cell qPCR, it is applicable for single cell RNA-seq data since there the whole genome is measured. For instance, it has been shown that the covariance of cell cycle related genes can be used to correct for specific gene expression during cell cycle phases [Buettner *et al*., 2015]. However, this is not the focus of this work where we introduce a correction scheme that is based on a global characteristic of each sample, namely the cells’ volume, rather than on the correlations among the expression of different genes.

## 2 Methods

### The cell growth correction probability

Measuring the abundance of a particular mRNA in a single cell during its cell cycle yields a discrete transcript number *m*, which is generally greater than the transcript number *m*_0_ that we would find at the beginning of the cell’s cell cycle (*τ*=0). During cell cycle the size of the cell increases from its initial volume *V*_0_=*V*(*τ*=0) (at the beginning of its cell cycle) to *V*(*τ*>0). Cell cycle and cell growth are intimately related [Mir *et al*., 2011, Kafri *et al*., 2013] and the number of mRNA transcripts within the cell increases as the cell volume increases. Therefore, we assume the concentration of mRNA transcripts *m*/*V* to remain constant during cell cycle. To render the numbers of mRNA transcripts from single cells with different cell sizes comparable, we introduce the volume-dependent cell growth correction probability 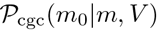. This is the probability of finding *m*_0_ mRNA transcripts within a cell’s initial volume *V*_0_ given a measured number of mRNA transcripts m within a cell’s total volume *V*. The volume-dependent cell growth correction probability is described by a binomial distribution

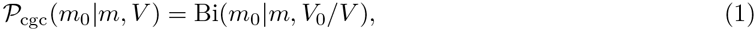

since this is the discrete probability distribution for finding *m*_0_ transcripts inside the initial volume *V*_0_ given the number of transcripts *m* present in the total volume *V* with success rate *p*=*V*_0_/*V* (see Figure 2A). In the limit of high mRNA transcript numbers the binomial distribution tends to a normal distribution. In this limit cell growth correction corresponds to scaling the measured number of mRNA transcripts *m* with the normalized volume of the cell *V*_0_/*V*. Therefore, the volume-dependent cell growth correction probability, Equation (1), contains the commonly performed scaling correction in the limit of high mRNA transcript numbers.

If the single cell’s volume *V* and its initial volume *V*_0_ are measured, we can evaluate 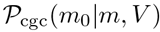 directly. In many experimental applications (such as qPCR), however, measuring each single cell’s volume is not performed or impossible. In this case, we treat the volume as a latent variable and marginalize over it to obtain the cell growth correction probability

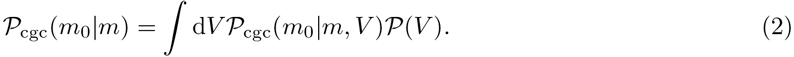

To evaluate this we require the probability distribution of the cells’ volumes 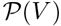 (i.e. the volume distribution over the cell population).This may be determined experimentally, or we can use generative models to simulate 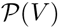 computationally. In the following we used a linear growth model to generate 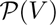 (see Supplementary Material S1). We evaluated the effect of different linear growth models in Supplementary Figure S2. The cell growth correction probability 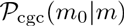 for linear growth is displayed in Figure 2B for several values of observed molecule numbers *m*.

**Fig. 2.**
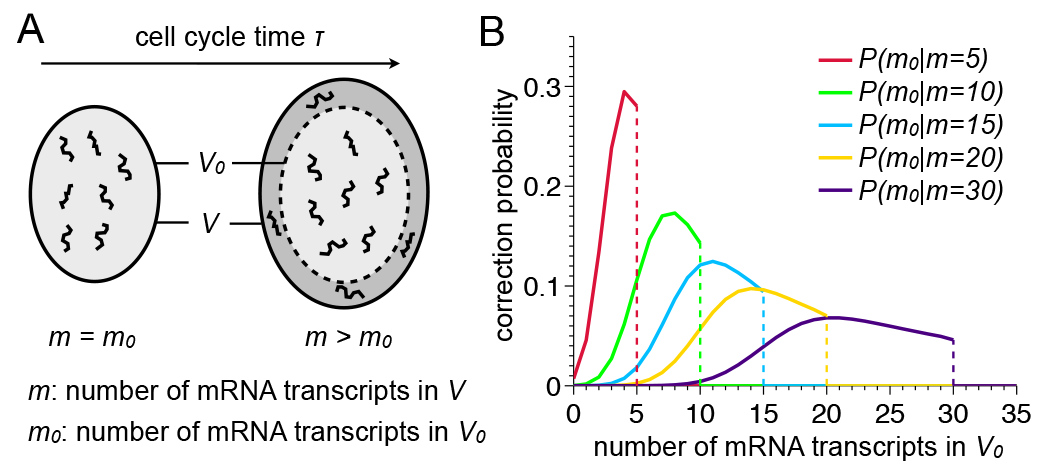
Cell growth model and correction probability. (A) During cell cycle, a cell will increase its initial volume *V*_0_ to *V*>*V*_0_. We assume the cell to increase its molecular content accordingly keeping the concentration of mRNA transcripts constant. Then the number of mRNA transcripts m measured at a latter time in cell cycle (*τ*>0) is greater than the number of mRNA transcripts *m*_0_ at the beginning of the cell cycle (*τ*=0). To render measured mRNA transcript numbers m from single cells at different time points in their cell cycle comparable, we calculate the cell growth correction probability 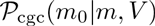. This is the probability to find *m*_0_ transcripts within the cell’s initial volume *V*_0_ (light grey area) given the cell’s current volume *V* and the measured number of transcripts *m*. (B) Cell growth correction probability 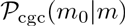 obtained after marginalizing over a linear growth-model (see text and Supplementary Figure S1) for several values of measured mRNA transcript numbers *m*. Notice that 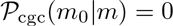 for *m*_0_>*m* resulting in the displayed discontinuities.

### cgCorrect for data normalization

The cell growth correction probability 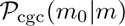 can be used to correct measured mRNA transcript numbers *m* directly to cell growth independent mRNA transcript numbers 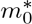 by determining its mode

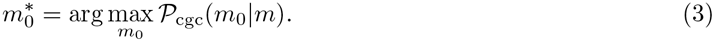

For instance, measuring *m*=15 transcript numbers in a single cell, the most likely value for the transcript number, which we corrected for differences in cell size is 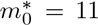 (see blue line in Figure 2B). This approach offers a rank-conserving, one-to-one correspondence between measured and cell growth corrected mRNA transcript numbers, as needed for normalization of a data set. When using point estimates (such as the mode of a probability distribution) many alternative mRNA transcript numbers *m*_0_ with non-negligible probability are ignored (see Figure 2B). However, we can also exploit the full distribution of the correction probability 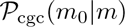 The number of mRNA transcripts of a particular gene is measured in many single cells. This yields a set of measured mRNA transcript numbers, which we use to obtain the steady-state probability distribution 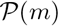 of measured mRNA transcript numbers of this gene. We then sum over the correction probability of all measured transcript numbers *m* multiplied by the steady-state probability distribution to gain the cell growth corrected steady-state distribution.

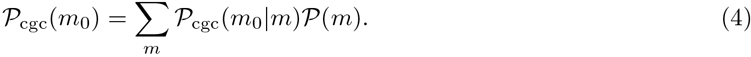

### cgCorrect for steady-state distribution analysis

The correction probability can also be used to account for differences in cell size when performing a steady-state distribution analysis of the transcript numbers of a particular gene. Given the mRNA transcript numbers *m* of a gene from several single cells, the likelihood 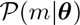 for the kinetic parameters ***θ*** of the underlying steady-state distribution can be calculated [Peccoud *et al*., 1995, Raj *et al*., 2006, Shahrezaei *et al*., 2008] (see Supplementary Material S2 for a summary of the analytical steady-state distributions for the simple and bursty gene expression mechanism). Neglecting differences in cell size, however, can lead to incorrect identification of the underlying steady-state distribution and its kinetic parameters (as already demonstrated in Figure 1C) and has not been considered within this context so far.

Using the correction probability, it is straightforward to incorporate cell growth correction into the existing framework of commonly performed steady-state distribution analysis,

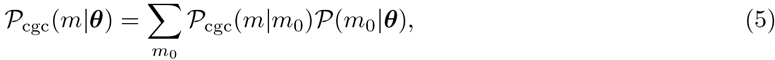

allowing us to obtain the likelihood for the measured mRNA transcript numbers *m* from cells that differ in cell size given the parameters ***θ*** of the steady-state distribution under consideration. To obtain 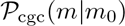 from the correction probability 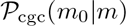, we use Bayes’ theorem with uniform prior on the measured transcript numbers numbers *m*.

For the simple gene expression mechanism the steady-state distribution is given by a Poisson distribution with one kinetic parameter: The effective transcription rate λ = *ν*/γ, which corresponds to the mean transcript number among all cells. For bursty gene expression the steady-state distribution is given by a negative binomial distribution with two kinetic parameters that allow for overdispersion: The burst size ξ =*ν*/*k*_off_ and the burst frequency *κ*_on_ = *k*_on_/γ (see Figure 1 and Supplementary Material S2 for details on the kinetic parameters). The model parameters can then be found via maximum likelihood estimation (MLE) 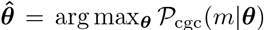. We evaluate if the parameters of both steady-state distributions are identifiable and therefore capable of describing the data by calculating their profile likelihoods [Raue *et al*., 2009].

If both the parameters of both distributions are identifiable, we perform model selection using the Bayesian information criterion (BIC) [Jeffreys, 1961, Kass *et al*., 1995] to select between the Poisson and the negative binomial distribution. Model selection based on the Bayesian information criterion (BIC) provides a good trade-off between goodness of fit and model complexity: by penalizing models with more parameters it counteracts overfitting the data. In case this model selection is inconclusive (ΔBIC ≤ 10), we call the underlying steady-state distribution inconclusive due to model selection (see Supplementary Material S3 for details on parameter estimation and model selection). As the BIC does not take into account the technical noise level of the data, we only perform model selection for those genes for which the biological variation significantly exceeds technical noise.

### cgCorrect and technical noise correction

In general, cell growth correction can also be combined with technical noise correction. To incorporate technical noise correction into the likelihood, Equation (5), the technical noise has to be measured in the experiment (e.g. with external spike-in controls) and the probability distribution of the technical noise 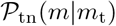 has to be determined experimentally. This is the conditional probability for the number of mRNA transcripts *m* that would be measured without technical noise given the number of mRNA transcripts *m_t_* that are measured and are subjected to technical noise. The likelihood of cell growth correction and technical noise correction can then be calculated as

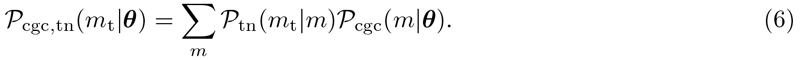

To obtain 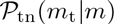 from the probability distribution of the technical noise 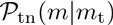, Bayes’ theorem can be applied with uniform prior on the measured mRNA transcript numbers with technical noise *m_t_*.

## 3 Results

### cgCorrect on simulated mRNA data leads to correct normalization and identification of the steady-state distribution

To validate cgCorrect, we applied it to mRNA transcript numbers that we simulated from the Poisson distribution (corresponding to the simple gene expression mechanism). We generated mRNA transcript numbers *m* of 100 single cells with different cell sizes (see Supplementary Figure S3 and Supplementary Material S4 for details on the simulation of the data).

Without differences in cell size the mRNA transcript numbers would be Poisson-distributed *m*_0_ ∼ Pois(*m*_0_|λ_o_) where the average number of mRNA transcripts per cell 〈*m*_0_〉=λ_o_ equals the effective transcription rate (see red line in Figure 1C and Figure 3A for an example where λ_0_=10). Due to differences in cell size the steady-state distribution of measured mRNA transcript numbers 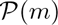 is shifted towards higher transcript numbers (green line in Figure 3A). We can correct for latent differences in cell size by calculating the corrected steady-state distribution of transcript numbers 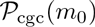, Equation (4) (blue line in Figure 3A). Since we ignored the cell’s volumes by marginalizing the volume out (cf. Equation (2)) the corrected steady-state distribution of transcript numbers does not entirely coincide with the Poisson distribution but has slightly larger tails. To compare cgCorrect with conventional house-keeping normalization, we scaled the measured number of transcripts *m* with the transcript number of an additionally simulated house-keeping gene *m_hk_* (see Supplementary Material S6 for details on house-keeping normalization), which we chose to have an average number of transcripts *m_0,hk_*=100 (yellow line in Figure 3A). Visual comparison of the two normalization strategies shows that cell growth correction for normalization outperforms house-keeping normalization for this data set.

Model selection based on steady-state distributions reports very strong evidence that the measured steady-state distribution of mRNA transcript numbers is can be rather described by the negative binomial than by the Poisson distribution and would therefore mistakenly be interpreted to origin from the bursty gene expression mechanism. When correcting for cell growth, model selection correctly identifies the steady-state distribution to be Poisson (see Supplementary Material S3 for details on parameter estimation and model selection). Performing parameter estimation the true effective transcription rate can only be inferred when using cgCorrect (see Figure 3B), confirming that cgCorrect outperforms house-keeping normalization in recovering the true underlying distribution. To test cgCorrect for a broad parameter range we simulated additional mRNA data sets for several average numbers of mRNA transcripts per cell (Figure 3C and D): Only when we apply cgCorrect we are able to infer the underlying steady-state distribution and its parameters for the whole parameter range correctly.

Moreover, we verified that after applying cgCorrect on transcript numbers that were simulated from the negative binomial distribution the inferred steady-state distribution is negative binomial: To this end, we simulated mRNA transcript numbers from the negative binomial distribution 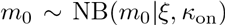 for a wide range of average numbers of mRNA transcripts (〈*m*_0_〉 = *ξ*. *k*_on_. When applying cgCorrect we found that model selection for *m*_0_〉 ≥ 3 correctly identifies the underlying steady-state distribution to be negative binomial. For very small average numbers of mRNA transcripts 〈*m*_0_〉 ≤ 2 the obtained distribution of transcript numbers is very narrow and we find cases (20% for 〈*m*_0_〉 = 2 and 90% for 〈*m*_0_〉 = 1), where the underlying steady-state distribution is identified to be Poisson (see Supplementary Figure S4). In summary, cgCorrect is capable of both, successfully inferring the underlying system parameters from the simulated, cell growth dependent transcript numbers and correctly specifying the steady-state distribution of transcript numbers.

When analyzing the steady-state distributions of genes one typically assumes that all cells of a particular cell type share the same kinetic parameters. This assumption does not necessarily reflect biological reality. To explore the effect of neglecting this assumption, we performed simulations where we varied the effective transcription rate of a gene simulated from the simple gene expression mechanism among all cells (see Supplementary Figure S5). Since the cells effective transcription rates differ among each other the simulated steady-state distribution may exhibit overdispersion and model selection may identify the steady-state distributions as being negative binomial.

**Fig. 3.**
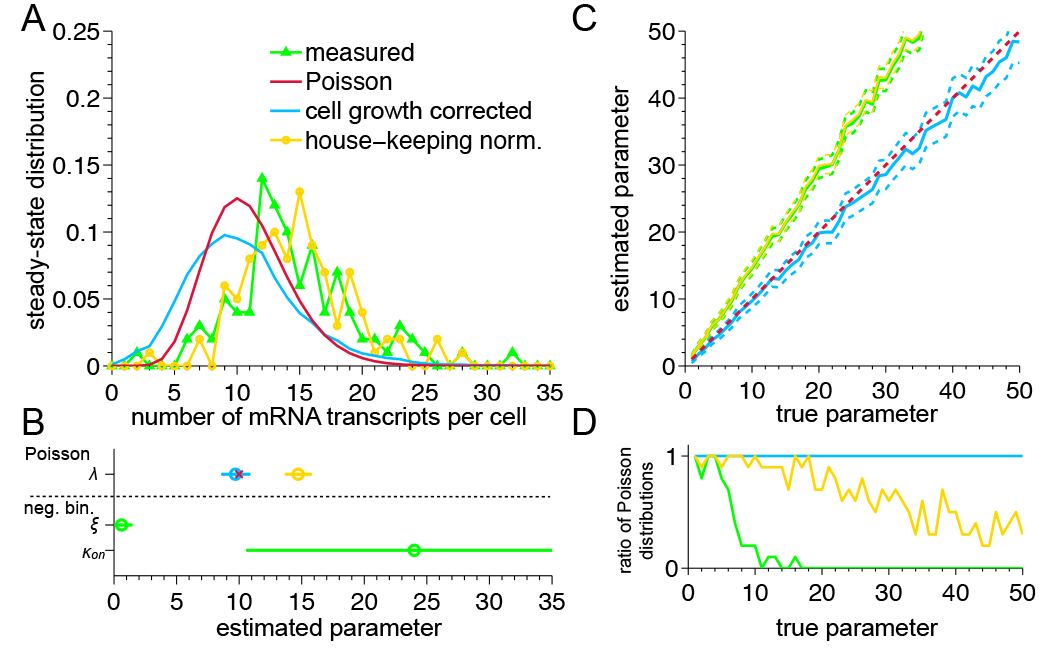
Cell growth correction of simulated gene expression data leads to the correct identification of parameters and the underlying steady-state distribution. (A) Steady-state probability distribution of the measured mRNA transcript numbers *m* (green line) and the cell growth corrected mRNA transcript numbers *m*_0_ (blue line). The underlying steady-state distribution of transcript numbers in the absence of differences in cell size is given by a Poisson distribution with an effective transcription rate λ_o_=10 (red line). The cell growth corrected probability distribution resembles the Poisson distribution closer than the house-keeping normalized distribution (yellow line). (B) Estimated parameters and identified steady-state distributions for cell growth corrected, house-keeping normalized and measured mRNA transcript numbers. For house-keeping normalization and cell growth correction the steady-state distribution is best described by the Poisson distribution and therefore their gene expression mechanism would be interpreted to be simple; for the measured data the steady-state distribution is best described by the negative binomial distribution and therefore the gene expression mechanism would be interpreted to be bursty. The Poisson distribution is governed by one parameter: the effective transcription rate λ_0_ = *ν*/γ. The negative binomial distribution is governed by two parameters: the burst size *ξ* = *ν*/*k*_off_ and the burst frequency *κ*_on_ = *k*_on_/γ. Only cell growth correction is in the range of the true parameter (red x). Error bars indicate 0.99 confidence intervals of the estimated parameters. (C) To explore the parameter range we performed parameter estimation and model selection for several values of the effective transcription rate λ_0_. We find that cell growth correction (blue line) is capable of correctly inferring the true parameter (red dashed line) for the whole parameter range whereas inferring the parameter on the measured (green line) and the house-keeping normalized mRNA transcript numbers (yellow line) fails. (D) Ratio of steady-state distributions that were identified to be Poisson for 10 independently simulated data sets. Model selection (based on the BIC) on the cell growth corrected data (blue line) identifies the true steady-state distribution correctly over the whole parameter range, in contrast to model selection on the measured data (green line) and on the house-keeping normalized data (yellow line).

**Fig. 4.**
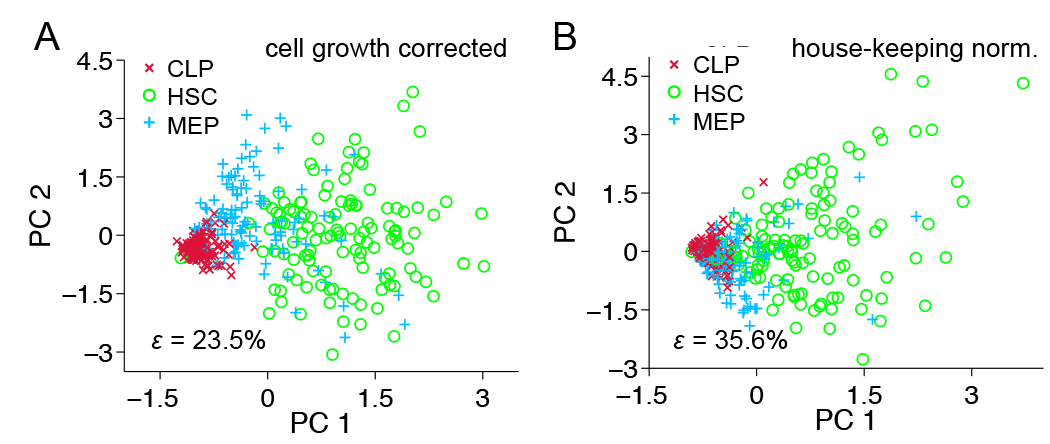
Principal component analysis (PCA) of single-cell qPCR data resolves hematopoietic sub-populations better when using cell growth correction. (A) PCA of cell growth corrected and (B) PCA of house-keeping normalized single-cell qPCR data of 18 transcripts. The nearest neighbor error ∈ decreases by 12.1% when using cell growth correction compared to house-keeping normalization.

### cgCorrect on qPCR data facilitates identifying distinct cell types and alters the interpretation of the gene expression mechanism based on a steady-state distribution analysis

To demonstrate the applicability of cgCorrect on single-cell qPCR data, we applied cgCorrect to a recently published data set of hematopoietic stem and progenitor (HSP) cells [Moignard *et al*., 2013]. In this experiment 18 transcripts of key hematopoietic genes (and six additional transcripts of house-keeping genes) were measured in 597 single cells of five different HSP cell types. To transform the measured data from ct-values into discrete numbers of mRNA transcript we use results from digital qPCR [Warren *et al*., 2006], where the discrete number of one of the 18 transcripts, *PU.1,* was measured for hematopoietic stem cells (HSCs), common lymphoid progenitors (CLPs) and common myeloid progenitors (CMPs), all of them found among the HSPs (see Supplementary MaterialS6 for details on the data pre-processing).

Since in this experiment neither technical noise nor information about the cells’ volume was measured we apply cgCorrect without technical noise correction and with marginalized volume (Equation 2). To compare cgCorrect with conventional house-keeping normalization we normalized the data set with the house-keeping genes *Ubc* and *Polr2a* as described by [Moignard *et al*., 2013]. cgCorrect is better suitable to resolve distinct cell types than house-keeping normalization, as can be visualized by a principal component analysis (PCA) (see Figure 4): The nearest neighbor error of finding two differing cell-types next to each other is decreased by 12.1%.

To further illustrate the effect of cgCorrect, we focus on one particular transcript (*PU.1*) in one cell type (CLP) (see Figure 5A). We analyze the Fano factor 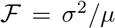 defined as the ratio between the variance σ^2^ and the mean µ of the steady-state distribution of mRNA transcript numbers. The Fano factor is a key parameter to quantify deviations from a Poisson distribution [Munsky *et al*., 2012] and it equals 1 if the values are Poisson-distributed. cgCorrect alters the Fano factor from 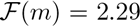 for the measured *PU.1* transcript numbers to 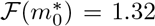. Parameter estimation for the measured and the corrected transcript numbers is depicted in Figure 5B. cgCorrect alters the identified steady-state distribution of *PU.1* in CLPs from following the overdispersed negative binomial distribution (in case of no correction) to Poisson.

**Fig. 5.**
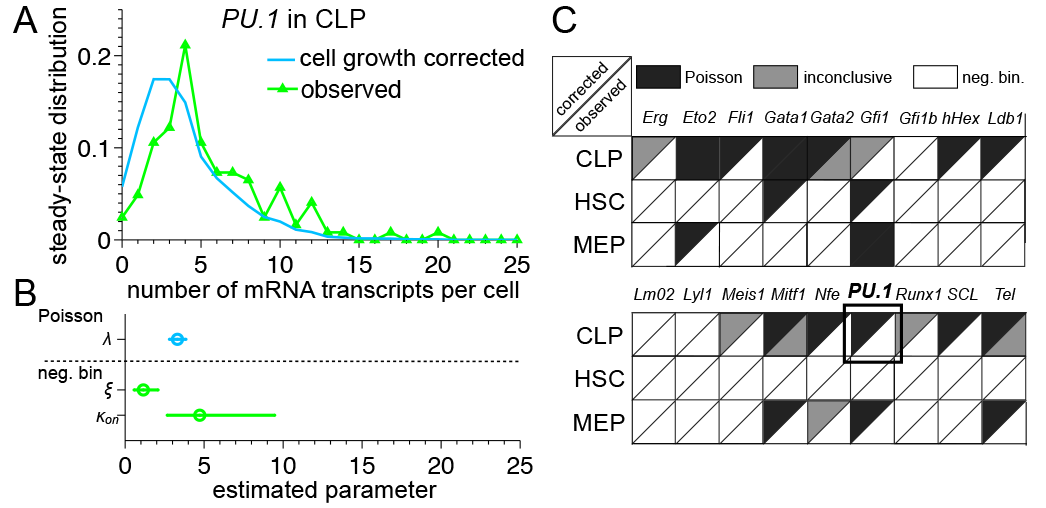
Parameter estimation and model selection on cell growth corrected steady-state distributions from single-cell qPCR data alters the interpretation of 15 out of the 56 hematopoiesis genes to origin more likely from the simple than from the bursty gene expression mechanism. (A) Probability density of corrected (blue) and measured (green) *PU.1* transcript numbers within CLPs. cgCorrect renders the observed, overdispersed distribution narrower.(B) Estimated kinetic parameters and identified steady-state distribution for *PU.1* mRNA in CLPs. The steady-state distribution is identified to be Poisson after cell growth correction (blue) and negative binomial without cell growth correction (green). cgCorrect alters the interpretation of the underlying gene expression mechanism from bursty to simple. (C) The result of model selection between the Poisson (black) and the negative binomial distribution (white) is visualized. Inconclusive model selection is indicated in grey. The lower right triangle indicates the identified steady-state distribution of the measured transcript numbers, the upper left triangle of the corrected transcript numbers. After performed cgCorrect the identified steady-state distribution is altered in 20 cases.

Applying cgCorrect to all measured mRNA transcripts, we find that the steady-state distributions of 18 out of 54 (∼ 33.0%) gene/cell type combinations are identified to be Poisson and would be interpreted to origin from the simple rather than the bursty gene expression mechanism, whereas this is the case for only 3 out of 54 (∼ 5.6%) without cgCorrect (see Figure 5C). A corresponding analysis with house-keeping normalization yields that the steady-state distribution of only 2 out of 54 gene/cell type combinations follow the Poisson distribution (see Supplementary Figure S6).

## 4 Discussion

In this work we present cgCorrect, a statistical method for the correction of latent differences in cell size. We show that differences in cell size may lead to an overdispersed steady-state distribution of transcript numbers, which may be misleadingly interpreted in a computational analysis. cgCorrect can be used for data normalization before visualization as well as for a steady-state distribution analysis of the data. It can incorporate information about the cell size on different levels: (i) If the size of each cell or an estimator for the size is known, we can use this information to obtain the volume-dependent cell growth correction probability. (ii) If only the probability distribution of the cells’ volume among the whole population is known we can use this distribution to marginalize the volume out. (iii) If there is a total lack of information about the cells’ volume (as is typically the case for qPCR data including the qPCR data set we analyzed), we can use generative growth models to simulate the cells’ volume distribution computationally and use this for marginalization.

We validated cgCorrect on simulated mRNA data, where we could show that it is only possible to infer the true steady-state distribution and its parameters when cgCorrect was applied. Analyzing steady-state distributions of transcript numbers from a qPCR data set we found that cgCorrect changed the identified steady-state distribution in 27.4% of the measured cell/gene combinations in HSPs from an overdispersed negative binomial distribution to the Poisson distribution.

In contrast to conventional normalization techniques cgCorrect takes the discreteness of mRNA transcript numbers into account. For the analyzed qPCR data set we showed that cgCorrect outperforms traditional house-keeping gene normalization resulting in a better separation of known cell types in a principal component analysis (PCA). House-keeping genes underlie stochastic gene expression themselves and may therefore not suit as reliable reporters for cell size.

In previous analysis the steady-state distribution of a gene is used to interpret its gene expression mechanism [Raj *et al*., 2006, Shahrezaei *et al*., 2008, Larson, 2011, Kim *et al*., 2013]. The Poisson steady-state distribution corresponds to the simple gene expression mechanism and the negative binomial distribution corresponds to the bursty gene expression mechanism. However, there are several assumptions involved that are important to consider for this interpretation.

First, it is assumed that the reaction rates that govern the gene expression mechanism remain constant during cell cycle. Here, we do not consider transcriptional changes during cell cycle that may alter the reaction rates and have been reported to affect the measured number of mRNA transcripts [Bertoli *et al*.,2013, Zopf *et al*., 2013]. In order to assess the effect of cell cycle specific gene expression, we modeled transcriptional changes of the reaction rates by an activation function reaching its maximum in the S phase of cell cycle. The resulting steady-state distribution is identified to follow the overdispersed, negative binomial distribution and would therefore be interpreted to origin from the bursty gene expression mechanism both with and without applying cgCorrect (see Supplementary Figure S7). Cell cycle specific gene expression corresponds to a highly orchestrated on and off switching of the promoter region. For a sample of unsynchronized cells that are pooled together, however, the resulting steady-state distribution of mRNA transcript numbers exhibits overdispersion.

The second assumption that is made when analyzing steady-state distribution of mRNA transcript numbers is that the kinetic parameters that govern gene expression are equal for all cells of the same cell type [Thattai *et al*., 2001, Shahrezaei *et al*., 2008, Raj *et al*., 2008, Kim *et al*., 2013], which does not necessarily have to reflect the biological reality. We tested the effect on the steady-state distribution analysis when neglecting this assumption by simulating mRNA transcript numbers from a cell population with varying transcription rates expressing mRNAs with the simple mechanism and showed that this effect can also lead to overdispersed steady-state distributions (see Supplementary Figure S5). A final conclusion on the gene expression mechanism cannot be made based on steady-state distributions of gene expression alone but needs techniques that allow for spatial and temporal resolution such as fluorescence in situ hybridization (FiSH) [Raj *et al*., 2006, Hocine *et al*., 2012, Battich *et al*., 2013].

Finally, we made assumptions concerning the cell growth parameters for the generative growth model that we used to obtain the correction probability. The question whether mammalian cells grow linearly or exponentially is still under debate [Cooper, 2004, Popescu *et al*., 2014]. Here, we used a linear growth model, which has been reported to be appropriate for rat Schwann cells [Conlon *et al*., 2003] to computationally simulate the distribution of cell volumes. Moreover, we performed a sensitivity analysis (see Supplementary Figure S2) that investigates the effect of different linear cell growth scenarios on the correction probability and indicates that our findings are robust with respect to the growth scenario. As already discussed, cgCorrect does not rely on a generative growth model as it allows to include additional information on either each single cell’s volume or the distribution of the cells’ volume, if they are measured.

To summarize, we identified differences in cell size of proliferating cells to be a latent cause of confounding variability. We introduced cgCorrect, a statistical method that is capable to correct for this confounding cell cycle effect in gene expression data, which can be used for data normalization, parameter estimation and model selection. We validated cgCorrect on a simulated data set and applied it to single-cell qPCR gene expression data [Moignard *et al*., 2013] from mouse HSPs.

## Acknowledgement

The authors thank John Marioni and Jan Hasenauer for helpful discussions.
This work was supported by the Helmholtz Alliance on Systems Biology (project CoReNe), the European Research Council (starting grant LatentCauses), the Deutsche Forschungsgemeinschaft (SPP 1356 Pluripotency and Cellular Reprogramming) and the Studienstiftung des deutschen Volkes [T.B.].

